# Ciliogenesis associated kinase 1 accumulates in its inactive form during polycystic kidney disease progression

**DOI:** 10.64898/2026.05.13.724873

**Authors:** Alice Serafin, Charlène Coquil, Aurore Dupuy, Mattias F. Lindberg, Darren P. Wallace, Pamela V. Tran, Oxana Ibraghimov-Beskrovnaya, Yannick Le Meur, Emilie Cornec-Le Gall, Christelle Ratajczak, Laurent Meijer, Vincent J. Guen

## Abstract

Ciliogenesis associated kinase 1 (CILK1) deficiency in human and mice results in kidney developmental defects including cystogenesis. However, the biology of CILK1 in autosomal dominant polycystic kidney disease (ADPKD), the most common inherited kidney disease, remains to be investigated. Here, we show that *CILK1* is overexpressed in dedifferentiated cells of renal tissue from ADPKD human patients in comparison to normal control tissue samples. We demonstrate that *CILK1* overexpression results in protein accumulation in a non-phosphorylated inactive form. Using mouse polycystic kidney disease models, we reveal that inactive CILK1 accumulation is progressive over the course of disease progression. We show that genetic inactivation of the *Polycystic Kidney Disease 1* (*PKD1*) gene is sufficient to trigger CILK1 accumulation. Altogether, these findings demonstrate that CILK1 regulation is altered in ADPKD and it represents a hallmark of disease progression.

## Introduction

ADPKD affects more than 10 million people worldwide, constituting a major public health burden (1). It represents the most common genetic renal disease, which is characterized by the development of renal cysts causing end-stage renal failure in adulthood (1). Pathogenic mutations in *PKD1* or *PKD2* genes are the most common cause of ADPKD (1,2). One mutant allele of *PKD* genes is generally inherited through germline transmission and the second mutant allele appears in a somatic manner leading to disease onset (3). Mutations in *PKD1* are responsible for almost 80% of cases and 15% of cases are attributed to mutations in *PKD2* (1,2). The remaining 5% of ADPKD cases are linked to rare mutations in other loci (e.g. *PKHD1 and PKHD2, HNF1B, GANAB, DNAJB11, SEC63, PRKCSH*), causing ADPKD-like phenotypes, or are of unclear genetic origin (1,2,4).

*PKD1* and *PKD2* genes encode polycystin-1 and -2 (PC1, PC2), respectively (1,3). PC1 and PC2 are transmembrane proteins (1,3). Pathogenic mutations of *PKD1* and *PKD2* result in loss of polycystins function (3). These transmembrane proteins interact with one another and form a membrane-bound receptor/ion channel complex (5,6). The N-terminal domains of PC1 protein is suspected to act as a mechanosensor and the C-terminal intracellular domains of PC1 and PC2 act in cell signaling (3,7-9). Polycystins are expressed at the highest level during pre-natal development of the kidney and their expression levels decreases in the adult tissue (10). In kidney epithelial cells, polycystins localize to different subcellular compartments and most notably to the primary cilium (3,11). Primary cilia are microtubule-based structures that act as a signaling hub at the apical cell surface (12). At the ciliary membrane, polycystins act as calcium channels (3). Upon polycystins loss of function, dysfunctional primary cilia promote cystogenesis (3,13). The precise ciliary signaling defects contributing to cystogenesis in ADPKD remain incompletely understood (3). In this study, we investigated CILK1 regulation in ADPKD.

CILK1 is a serine/threonine protein kinase that localizes to primary cilia whose deficiency in human leads to skeletal, endocrine, central nervous system and kidney disorders (14-18). In mice, genetic inactivation of *CILK1* results in kidney defects associated with cystogenesis (19). The role of CILK1 in development is suspected to be mediated by its function in ciliary signaling. CILK1 kinase activity controls proper movement of protein complexes along the ciliary axoneme by regulating intraflagellar transport (15,20). CILK1 regulation of cilium elongation relies on its kinase activity and phosphorylation on different residues (18,21). FGFR kinases phosphorylate CILK1, thereby repressing its kinase activity, and CDK20 phosphorylates it, thereby promoting its activation (18). CILK1 kinase activity also relies on autophosphorylation of Y159 for its basal activity (21). However, CILK1 regulation in ADPKD remains poorly understood.

Here, we assessed expression of *CILK1* as well as the abundance and phosphorylation of its protein product in different forms of PKD. We demonstrate that *CILK1* gene is overexpressed in multiple kidney segments of human ADPKD renal samples in comparison to control tissue samples. *CILK1* overexpression results in increased abundance of the protein, which accumulates in a Y159 non-phosphorylated form, representing inactive CILK1. Using mouse PKD tissue samples, we show that inactive CILK1 accumulates in different models of cystogenesis and this accumulation is progressive over the course of disease progression. We demonstrate that genetic inactivation of *PKD1*, the main cause of ADPKD, is sufficient to trigger CILK1 accumulation in renal cells. Altogether, our findings establish CILK1 abnormal regulation as an important feature of ADPKD. Our data reveal mechanistic insights into abnormal regulation of a critical ciliary signaling component in ADPKD.

## Results

### *CILK1* is overexpressed in different segments of ADPKD renal tissues

We initiated this study by analyzing gene expression in cells of human ADPKD renal tissue samples and in cells of non-tumoral renal samples of cancer patients, as controls, using public single-cell RNA sequencing data (Figure 1A) (22). 118 561 nuclei (controls: 42 217, ADPKD: 76 344) were analyzed after quality control. We conducted unsupervised clustering, based on differential gene expression, which yielded twelve different cell type clusters reflecting the cellular diversity in the kidney (Figure 1B-C). Using canonical lineage markers, we detected the cellular constituents of each cluster (Figure 1D, including: nephron epithelial cells, fibroblasts (FIB: *ACTA2+, COL12A1+, COL1A1+*), endothelial cells (ENDO: *EMCN+, FLT1+*), leucocytes (LEUK: *PTPRC+, CTSD+*) and urothelial cells (URO: *UPK1A+, UPK3A+*). Among nephron epithelial cells, we distinguished podocytes (PODO: *NPHS2+, PODXL+*), parietal epithelial cells (PEC: *ALDH1A2+, CFH+*), proximal cells (PT: *MOX+, SLC17A1+, SLC13A3+*), thick ascending loop of Henle cells (TAL: *SLC12A1+, UMOD+, GP+*), distal connecting tubule cells (DCT: *SLC12A3+, TRPM6+, KILH3+*) and collecting duct cells, including principal cells (PC: *SCNN1G+, FXYD4+, AQP2+*), intercalated A (ICA: *DMRT2+, SLC26A7+, ADGRF5+*) and B (ICB: *INSRR+, SLC26A4+, SLC4A9+*) cells. We detected all cell types in both control and ADPKD tissue samples except LEUK and URO that derived exclusively from ADPKD samples (Figure S1A). Major transcriptional changes were apparent between cells from control and ADPKD tissue samples within individual clusters (Figure 1C), reflecting general cellular de-differentiation in the disease state associated with downregulation of expression of canonical cell type markers (Figure 1D).

**Figure 1.**
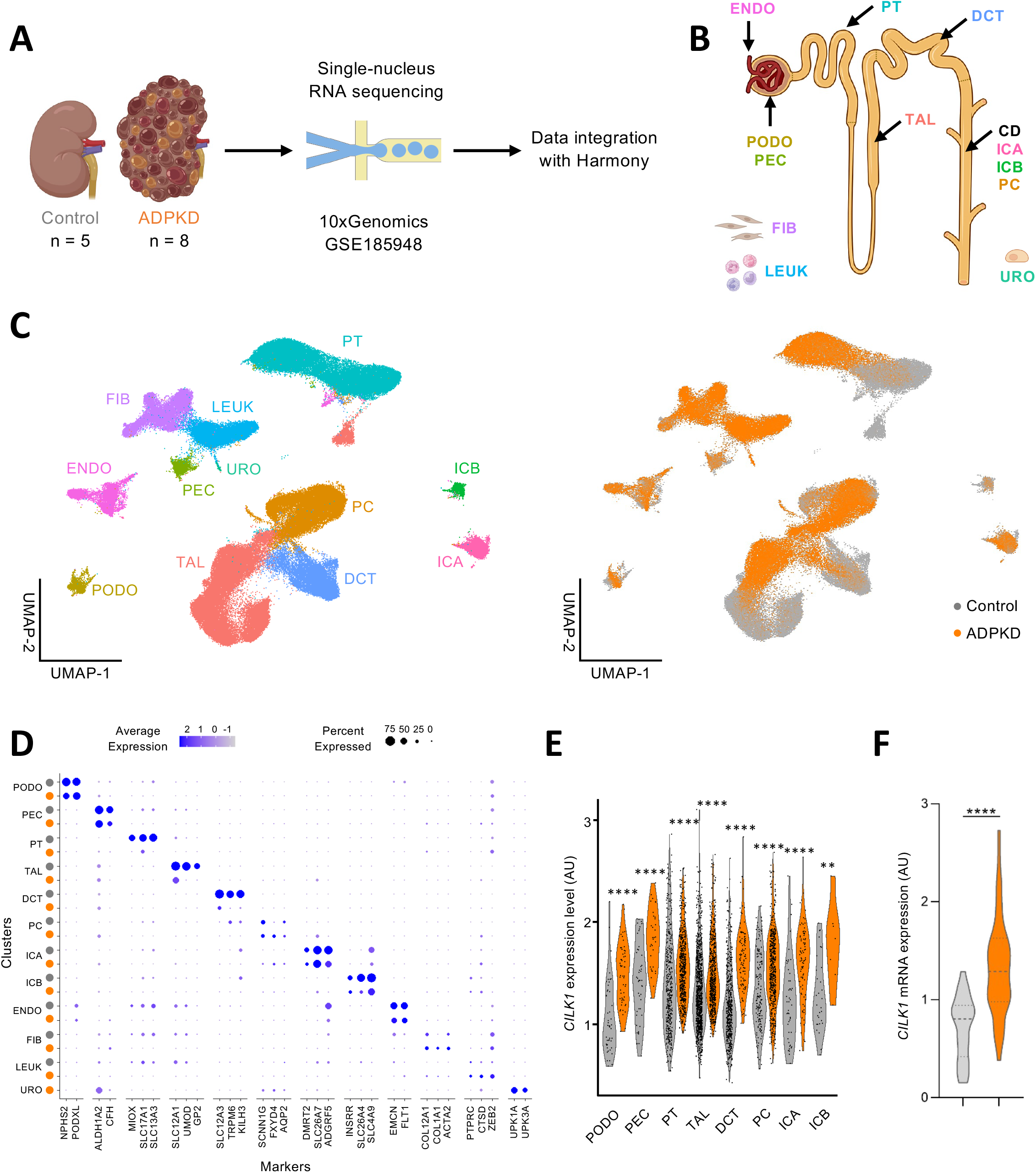
*CILK1* upregulation in all tubular cell types in ADPKD. (A) Overview of experimental methodology used by Muto et al. (22): 5 control and 8 ADPKD kidney samples were analyzed with snRNA-seq and data were integrated with Harmony. (B) Overview of kidney cell types: PODO podocytes; PEC parietal epithelial cells; PT proximal tubule; TAL thick ascending limb of Henle’s loop; DCT distal convoluted tubule; CD collecting duct; PC principal cells; ICA Type A intercalated cells; ICB Type B intercalated cells; ENDO endothelial cells; FIB fibroblasts; LEUK leukocytes; URO uroepithelium. (C) UMAP plot of integrated snRNA-seq dataset with annotation by cell type (left) or disease condition (right). (D) Dot plot of snRNA-seq dataset showing gene expression patterns of cluster-enriched markers for control (grey) and ADPKD (orange) kidneys. Only data from ADPKD kidneys were shown for URO cluster. Dot diameter corresponds to the proportion of cells expressing the indicated gene and dot intensity corresponds to average expression relative to all cell types. (E) *CILK1* expression level depending on kidney epithelial cell types for control (grey) and ADPKD (orange) kidneys. Violin plot (cut off expression level = 0.2). Each dot represents one cell. Wilcoxon test: \*\**P*<0.01, \*\*\*\**P*<0.0001. (F) *CILK1* mRNA expression in controls (n=8 from Brest cohort and n=14 from Kansas cohort) and ADPKD patients (n=35 *PKD1* and n=5 *PKD2* from Brest cohort, and n=12 *PKD1* from Kansas cohort). Violin plot. Unpaired Student’s t test: \*\*\*\**P*<0.0001.

Having defined the identity of the cell clusters forming the single-cell landscape, we compared *CILK1* expression between all nephron epithelial cells from ADPKD samples to those from control samples. We found that only a subpopulation of cells in both type of samples expressed *CILK1* (Figure S1B). Most notably, *CILK1*-expressing cells from ADPKD samples expressed significantly higher *CILK1* transcript levels compared to those from control samples (Figure S1C). Gene-set enrichment analysis revealed that *CILK1*^high^ ADPKD cells express higher levels of a gene regulatory network of apoptosis compared to *CILK1*^low^ and *CILK1-negative* cells (*P*=1.3×10^-10^ and *P*<2.2×10^-16^, respectively, Figure S1D-F, Supplementary table 1). To determine whether the subset of *CILK1*^high^ cells in ADPKD derive from a specific kidney segment, we next assessed *CILK1* levels in each of the 8 different nephron epithelial cell types. We repeatedly found higher transcript levels in cells from ADPKD samples compared to control samples for each kidney segment, supporting general *CILK1* overexpression across polycystic kidneys (Figure 1E). As an alternative approach, we assessed *CILK1* expression in control and human ADPKD renal tissue samples from two distinct patient cohorts using real-time qPCR (Brest cohort: n=40, Kansas City cohort: n=12). We found significantly higher *CILK1* transcript levels in ADPKD patient samples compared to controls (*P*<0.0001, Figure 1F, Figure S1G). Collectively, these data establish that *CILK1* overexpression is a general transcriptional hallmark of cells from different kidney segments in ADPKD patients.

### CILK1 protein accumulates in an inactive form in ADPKD

We reasoned that increased *CILK1* transcripts could translate into increased protein levels in ADPKD samples. To address this hypothesis, we conducted Western blotting using kidney protein extracts from our two distinct patient cohorts. We found significantly higher levels in CILK1 total protein levels in ADPKD samples harboring either *PKD1* or *PKD2* mutations compared to control samples (*P*<0.0001, Figure 2A-B, Figure S2A-C). Given the observation that CILK1 loss of function in a mouse model results in renal cystogenesis, we therefore asked whether the accumulation of CILK1 that we detected in ADPKD tissue samples reflects accumulation of inactive CILK1. CILK1 auto-phosphorylates on Y159 and this phosphorylation event reflects its kinase activity (21). Immunoblotting for Y159 phosphorylated CILK1 (pY159-CILK1) revealed significant lower levels of pY159-CILK1 in ADPKD samples compared to control samples (*P*<0.0001, Figure 2A-B, Figure S2D-F), supporting the notion that CILK1 accumulates in an inactive form in the renal tissue of ADPKD patients. To determine whether CILK1 accumulation can be detected through standard immunohistology, we performed CILK1 and pY159-CILK1 immunostaining of renal tissue sections from control and ADPKD patients (Nantes cohort). While pY159-CILK1 was below the detection limit, this analysis revealed significantly higher abundance of total CILK1 in ADPKD kidney sections both in tubular and peri-tubular areas compared to controls across kidney sections (*P*=0.0077, Figure 2C-D). Collectively, these data reveal that inactive CILK1 accumulation can be detected through different routine methods as a general hallmark of renal tissue degeneration in ADPKD.

**Figure 2.**
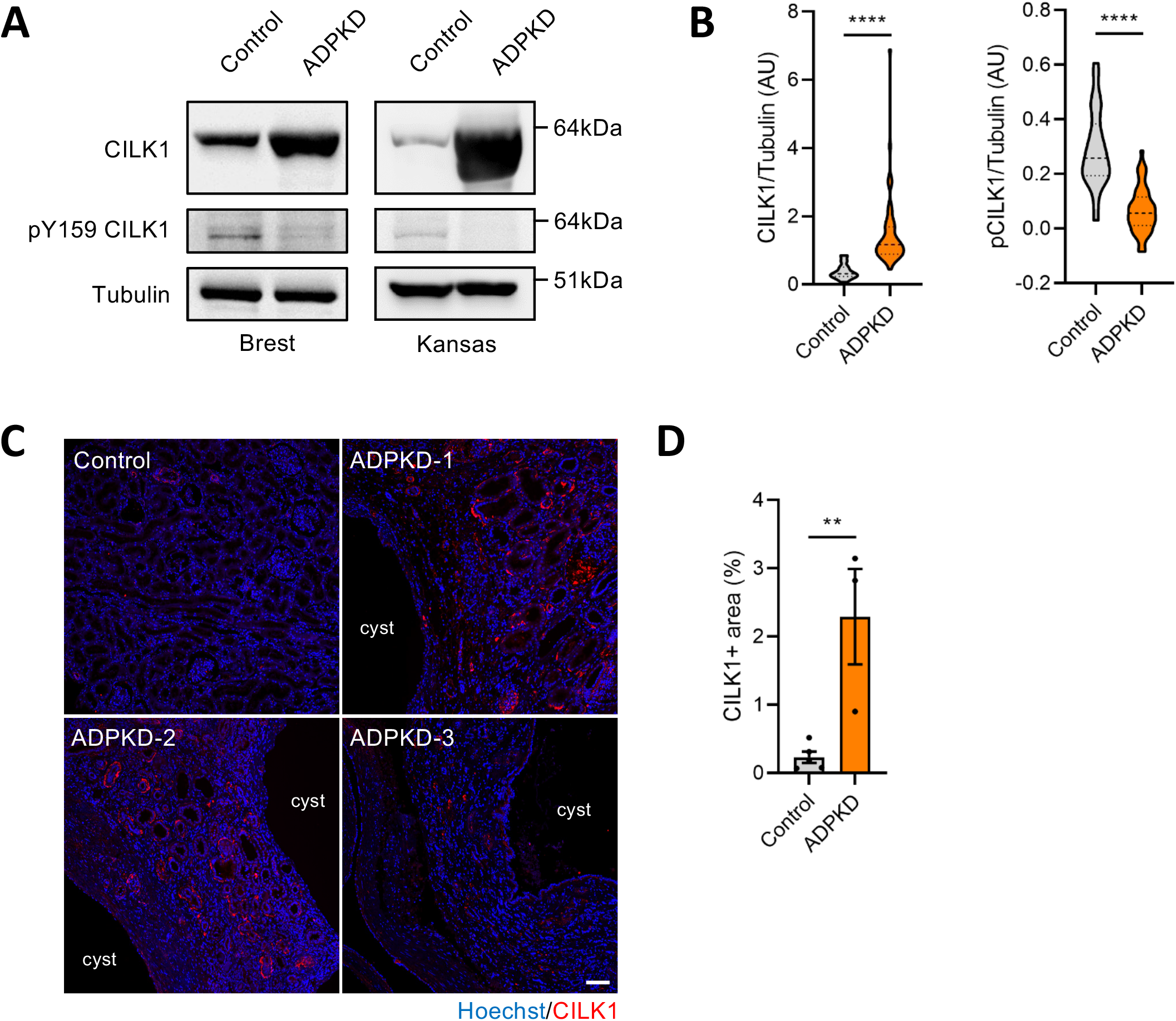
Increased CILK1 inactive form in ADPKD patients. (A) Representative images of Western blot analysis showing CILK1, p-CILK1 (Y159) and Tubulin in controls and ADPKD patients from Brest and Kansas cohort. (B) Quantification of the relative abundance of CILK1 (left) and p-CILK1 (right) in controls (n=8 from Brest cohort and n=14 from Kansas cohort) and ADPKD patients (n=35 *PKD1* and n=5 *PKD2* from Brest cohort, and n=12 *PKD1*). Violin plot. (C) Representative image of immunostaining of a control (commercially available samples) and 3 ADPKD patients (Nantes cohort) showing nuclei (blue) and CILK1 (red). Scale bar = 100µm. (D) Quantification of CILK1-positive staining area in kidney sections from controls (n=5) and ADPKD patients (n=3). Each dot represents one control/patient. Mean ± SEM. (B, D) Unpaired Student’s t test (B, p-CILK1 ; D) or Mann-Whitney test (B, CILK1): \*\**P*<0.01, \*\*\*\**P*<0.0001.

### Inactive CILK1 accumulates in the renal tissue of different murine PKD models and this is progressive over the course of disease progression

To determine whether CILK1 also accumulates in its inactive form in tissues of PKD models of different genetic origin, we examined tissue extracts from 8 different mouse models, each of which harboring a distinct genetic alteration but all resulting in ciliary signaling dysfunction and polycystic kidneys (Figure 3A, Table 1). These very same models had been used to investigate the renal targets of (R)-roscovitine and (S)-CR8, two drugs active in ADPKD models (23-25). Using Western blotting, we repeatedly found significantly higher levels of CILK1 total protein (*P*=0.0097) and lower levels of pY159-CILK1 in murine polycystic models (*P*=0.0019, Figure 3A-B), supporting the notion that impaired CILK1 regulation is a shared feature of different forms of polycystic kidney disease. To examine the dynamics of CILK1 modulation in PKD, we next assessed CILK1 and pY159-CILK1 levels in protein extracts from renal samples collected from *jck* mice at distinct stages of disease progression. In this model, cysts emerge in some kidney segments as early as post-natal day 3, further develop across different segments by day 50, and grow progressively until day 100 (26). We therefore examined CILK1 and pY159-CILK1 levels in renal tissue extracts over the first 100 post-natal days by Western blotting. This showed that by day 50, CILK1 protein levels starts to increase and pY159-CILK1 levels start to decrease (Figure 3C). We next detected progressive escalation of these phenotypes until day 100 (Figure 3C). Thus, we conclude that inactive CILK1 accumulation is progressive during kidney cystogenesis.

**Table 1.** Murine models used in this study.

| Model | Name | Mutation/ Deletion | Age/Disease stage |
| --- | --- | --- | --- |
| *Pkd1*<sub>TAG 26</sub> | Polycystic kidney disease 1 (polycystin 1) | Mutation on *Pkd1* gene in renal and extra renal tissues | 8 months / advanced disease |
| *Pkd1* extra 11 | Polycystic kidney disease 1 (polycystin 1) | Deletion of a fragment of the *Pkd1* gene. Expresses only extracellular domain of PKD1 | 19 months / advanced disease |
| *Pkd1*<sup>flox/-</sup>; Ksp-Cre | Polycystic kidney disease 1 (polycystin 1) | *Pkd1* gene inactivated in kidneys | Postnatal day 7 / very advanced disease |
| *Bpk* | BALB/c polycystic kidney | Abnormal 3' coding region of the *bicaudal C homolog 1* gene, *Bicc1* | Postnatal day 21 / very advanced disease |
| *Jck* | Juvenile cystic kidney | Missense mutation on *Nek8* gene, G448V, in the highly conserved RCC1 domain of NEK8 | 26, 39, 45, 50, 64 and 100 days / advanced disease from 45 days |
| *Cpk* | Congenital polycystic kidney | Mutation in the *Cys1* gene | 19 days / very advanced disease |
| *Pcy* | Polycystic | Missense mutation (T1841G) in the mouse orthologue of the human *Nphp3* gene (encoding Nephrocystin-3) | 15 weeks / moderate disease |
| *Pkd1*<sup>RC/RC</sup> | Polycystic kidney disease 1 (polycystin 1) | Knockin mouse model of a likely disease variant, *PKD1*p.R3277C (RC) on C57B6 background | 6 months / moderate disease |

**Figure 3.**
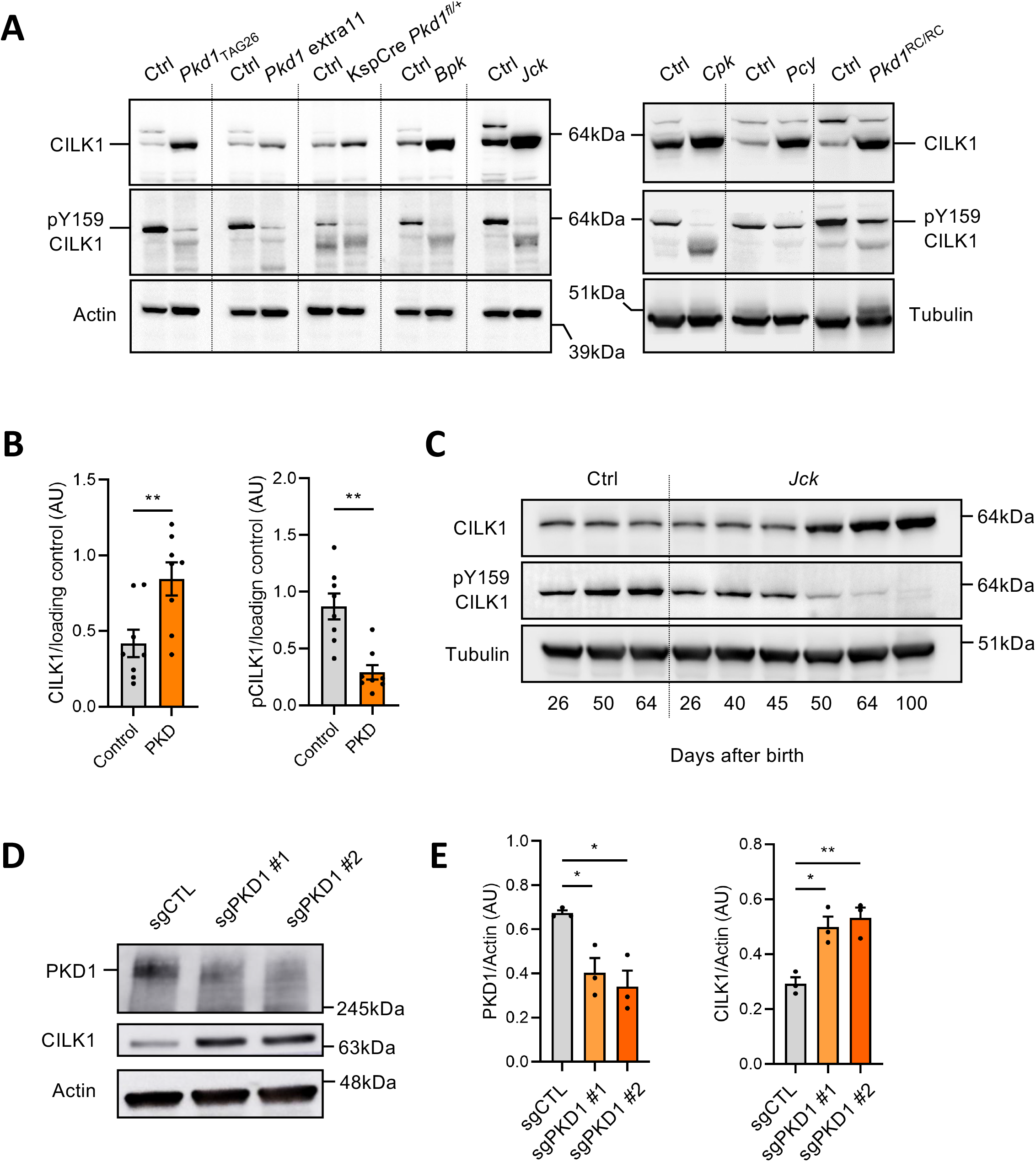
Increased CILK1 inactive form during polycystic kidney progression and after PKD1 genetic inactivation. (A) Western blot analysis showing CILK1, p-CILK1 (Y159) and Actin/Tubulin in control mice (Ctrl) and 8 polycystic murine models (Table 1). (B) Quantification of the relative abundance of CILK1 (left) and p-CILK1 (right) in controls and mice with polycystic kidney disease (PKD). Each dot represents one mouse (n=8 per group). (C) Western Blot analysis showing CILK1, p-CILK1 (Y159) and Tubulin over time in control and *Jck* mice. (D) Representative image of Western blot analysis showing PKD1, CILK1 and Actin in mIMCD3 control cells (sgCTL) and in two mIMCD3 polyclonal cells invalidated for PKD1 (sgPKD1). (E) Quantification of the relative abundance of PKD1 (left) and CILK1 (right) in mIMCD3 sgCTL and sgPKD1. Each dot represents one experiment (n = 3 experiments). (B, E) Mean ± SEM. Unpaired Student’s t test (B, CILK1 ; D) or Mann-Whitney test (B, p-CILK1): \**P*<0.05, \*\**P*<0.01.

### *PKD1* genetic inactivation is sufficient to alter CILK1 expression in renal epithelial cells

PKD in mouse models used to explore CILK1 regulation is caused by either direct genetic inactivation of *PKD1* genes or of genes that affect function of the *PKD1* gene product (27-29). Thus, we hypothesized that loss of function of *PKD1* is the critical event triggering CILK1 abnormal regulation. To test this hypothesis, we used a lentiviral CRISPR/Cas9 strategy to mutate *PKD1* in murine inner medullary collecting duct cells (mIMCD3) (30), and we immediately assessed the impact on PC1 using western blotting. This yielded cell populations with partial but significant reduction of PC1 protein levels for each of the guide RNA tested (sgPKD1#1 and #2) compared to a control guide RNA (sgCTL, *P*=0.0165 and *P*=0.0111, Figure 3D-E), due to the cell-to-cell variability in the inactivation of *PKD1*. Notably, we detected significant upregulation of CILK1 levels in response to *PKD1* mutation compared to wild-type controls (*P*=0.0102 and *P*=0.0056, Figure 3D-E). Together, these data show that genetic inactivation of *PKD1* is sufficient to trigger CILK1 accumulation in renal epithelial cells.

## Discussion

Autosomal polycystic kidney disease is one of the most common ciliopathies caused by loss of function of the polycystin ciliary proteins. However, the ciliary signaling defects resulting from deficiency of these components remain incompletely understood at the mechanistic level. Here, we examined regulation of the ciliary protein kinase CILK1 in renal tissue samples of ADPKD patients and using a combination of murine polycystic kidney disease models. Previous reports revealed that *CILK1* genetic inactivation causes cystic phenotypes in mice. Our data reveal transcriptional and post-transcriptional alterations of CILK1, resulting from polycystin loss of function, as a general hallmark of ADPKD in humans and mice. Thus, our findings further expand the spectrum of CILK1 alterations linked to cystogenesis, offering new insights into abnormal regulation of yet another ciliary component in ADPKD progression.

We detected accumulation of CILK1 in tubular and peri-tubular regions in ADPKD renal tissue sections, suggesting that CILK1 accumulates also outside primary cilia of renal cells in the disease state. CILK1 localization to primary cilia in renal cells is well documented but cytoplasmic localization and extracellular secretion of CILK1 remain poorly documented. However, expression of a fluorescently tagged version of CILK1 in renal cells revealed cytoplasmic localization (15). Collectively, these data suggest ciliary and extra-ciliary dysfunction of CILK1 in ADPKD progression.

We found that *CILK1*-overexpressing cells also co-express higher levels of a signature of genes acting as regulators of apoptosis. Regulated cell death in ADPKD remains poorly understood. Initial studies in the field reported increased apoptosis in the renal tissue of different ADPKD rodent models (31-34). However, independent studies either reported no significant changes or decreased apoptosis in the renal tissue of such models (31,35,36). Genetic and pharmacological strategies to perturb apoptosis also resulted in confusing phenotypes in ADPKD models (37,38). The complex role of apoptosis in ADPKD likely reflects implication of distinct cell death modalities occurring in distinct renal cell types at different stages of disease progression. Our data support the notion that *CILK1*^*high*^ cells in renal epithelial cells of advanced ADPKD upregulate apoptosis regulating genes. Driver genes of this signature represent anti-apoptotic genes. This suggests that CILK1 alteration occurs in cells with enhanced ability to resist to cell death that potentially contributes to disease progression and severity.

There is an emerging interest in targeting ciliogenesis or ciliary signaling for genetic renal disease treatment, but a lack of inhibitors that efficiently and selectively achieve this goal. Incidentally, polycystic kidney disease provides an opportunity to investigate pathogenic ciliary dysfunctions and identify novel actionable vulnerabilities. Our data suggest that inhibition of proteins that act to repress the ciliary kinase CILK1 in ADPKD could represent a valuable approach for ADPKD treatment. FGFR protein kinases phosphorylate CILK1 to inactivate it (39), and inhibitors of these kinases represent therefore a prototypical class of small-molecules of therapeutic interest for ADPKD. It is also interesting to note how increased expression and dephosphorylation of CILK1 mirrors the overexpression of CSNK1ε and the alteration of CSNK1α isoforms pattern in polycystic kidneys (25). Whether the CILK1 and CSNK1ε/CSNK1α are connected remains to be determined. Additionally, several cyclin-dependent kinases (CDK2, CDK5, CDK20) have been shown to regulate ciliary functions and their inhibition by (R)-roscovitine or (S)-CR8 attenuates cystogenesis in PKD models (23,24,40). CDK20 (previously known as CCRK) phosphorylates CILK1 on Thr157, leading to its activation and modulation of intraflagellar transport (41,42). A deeper understanding of the interactions between CDKs and CILK1 in a physiological setting and during cystogenesis represents another interesting opportunity to identify additional actionable vulnerabilities and thus more effective treatment for ADPKD.

## Methods

Cell culture, immunostaining, real-time qPCR, and snRNA-seq data processing are detailed in Supporting Information.

### Human tissue samples and animal models

Human tissues were provided by Dr Darren P. Wallace (University of Kansas Medical Center, Kansas City, USA), Dr. Yannick Le Meur (CHRU La Cavale Blanche, Brest, France), the Tumorothèque of Nantes CHU (France) or purchased at Tissue Array (#KDN242). The data corresponding to human tissues are present in Supplementary Table 2. For mRNA and protein experiment, tissues were snap-frozen and stored at -80°C until further use. For immunostaining, tissues were fixed with formol, embedded in paraffin and stored at room temperature until further use. Mouse tissues were provided by different contributors. Human tissue collection and mouse handling were approved by committees of the partner institutions. The study abides by the Declaration of Helsinki principles.

### Antibodies

For Western blotting we used Anti-CILK1 (Assaybiotech B8112, 1:500), anti-phospho-Tyr-159 CILK1 (Assaybiotech P12-1118, 1:500), anti-tubulin (Abcam ab56676, 1:1000) and anti-β-actin (Abcam ab8227, 1:1000). For immunostaining, we used anti-CILK1 (Kinexus NK075.5, 1:200) and Hoescht (Invitrogen H3570, 1:5000).

### Protein extraction and western blot experiments

Tissues were crushed in homogenization buffer (5mL/g of material; 25mM Mops (pH 7.2), 15mM EGTA, 15mM MgCl2, 60mM β-glycerophosphate, 15mM p-nitrophenylphosphate, 2mM dithiothreitol, 1mM sodium orthovanadate, 1mM sodium fluoride, 1mM phenylphosphate disodium and 1X Roche complete protease inhibitors) with Minilys homogenizer (Bertin Technologies). Homogenates were centrifuged for 10min at 21,000g at 4°C. A part of the supernatant was diluted in half in 2X loading buffer with 400mM of dithiothreitol then denaturated at 99°C for 3min and assess for the protein content (BioRad DC Protein Assay). Protein extraction from IMCD3 cells was conducted using standard procedures as described previously (1-3). Proteins and protein standard (Seablue from Thermo Fisher or ColorMixed from Solarbio) were separated by 10% NuPAGE pre-cast Bis-Tris polyacrylamide mini gel electrophoresis (Invitrogen) with MOPS-SDS running buffer. Proteins were transferred to 0.45µm nitrocellulose membranes with Trans-Blot Turbo System (BioRad). These were blocked with 5% low fat milk in Tris-buffered saline/Tween 20 and incubated overnight at 4°C with antibodies. Secondary antibodies conjugated to horseradish peroxidase (BioRad) were added to visualize the proteins using the Enhanced Chemiluminescence reaction. Protein quantification was performed on FIJI v.1.54p by measuring pixel mean gray value.

### Statistics and data availability

Statistical calculations and figure representations were performed with R v.4.3.2 or GraphPad Prism v8. The Shapiro–Wilk test was used to test the normality. Differences between two groups that do not meet the normal distribution were compared using two-tailed Mann– Whitney U test. If not possible or if the two groups meet the normal distribution, differences were compared using unpaired two-tailed Student’s t test. For single nuclei analysis, we used Wilcoxon test. *P* <0.05 was considered statistically significant. Statistical tests and exact sample sizes used to calculate statistical significance are stated in the figure legends. All data are contained within the manuscript.

## Acknowledgments

We are thankful to the following researchers for providing kidney samples of their animal models: Alessandra BOLETTA, Thomas WEIMBS, Nikolay BUKANOV, Christopher WARD, Michal MRUG, and Marie TRUDEL. We thank BioCore (US16 Inserm – UAR 3556 CNRS), including IBISA MicroPICell facility (Biogenouest), member of the national infrastructure France-Bioimaging supported by the French national research agency (ANR-10-INBS-04). Human ADPKD tissues were provided by the Kansas Polycystic Kidney Disease Research and Translation Core Center of the PKD-Research Resource Consortium (U54 DK126126). This work was supported by Conseil Régional de Bretagne/Feder (LM), iSITE NExT ANR-16-IDEX-0007 (VG) and Fondation Groupama (VG).

## Author contributions

A.S., C.C., L.M., V.J.G. designed research; A.S., C.C., A.D. performed research, M.F.L. D.P.W., P.V.T., O.I.B., Y.L.M., E.C.L, C.R. contributed new reagents/analytic tools, A.S., C.C., A.D., L.M., V.J.G. analyzed data. A.S., L.M., V.J.G wrote the paper.

## Supporting information

This article contains supporting information including references (22,43-47).

## Footnotes

ADPKD: autosomal dominant polycystic kidney disease; AU: arbitrary unit; CD: collecting duct; CILK1: Ciliogenesis associated kinase 1; DCT: distal convoluted tubule; ENDO endothelial cells; FIB fibroblasts ICA: Type A intercalated cells; ICB: Type B intercalated cells; LEUK: leukocytes; ns: non significant; PC: principal cells; PC1/2: polycystin-1/2; PEC: parietal epithelial cells; PKD: Polycystic Kidney Disease; PODO: podocytes; PT proximal tubule; TAL: thick ascending limb of Henle’s loop; URO: uroepithelium.

