## Supporting information for "Ciliogenesis associated kinase 1 accumulates in its inactive form during polycystic kidney disease progression"

**Supplementary figures**

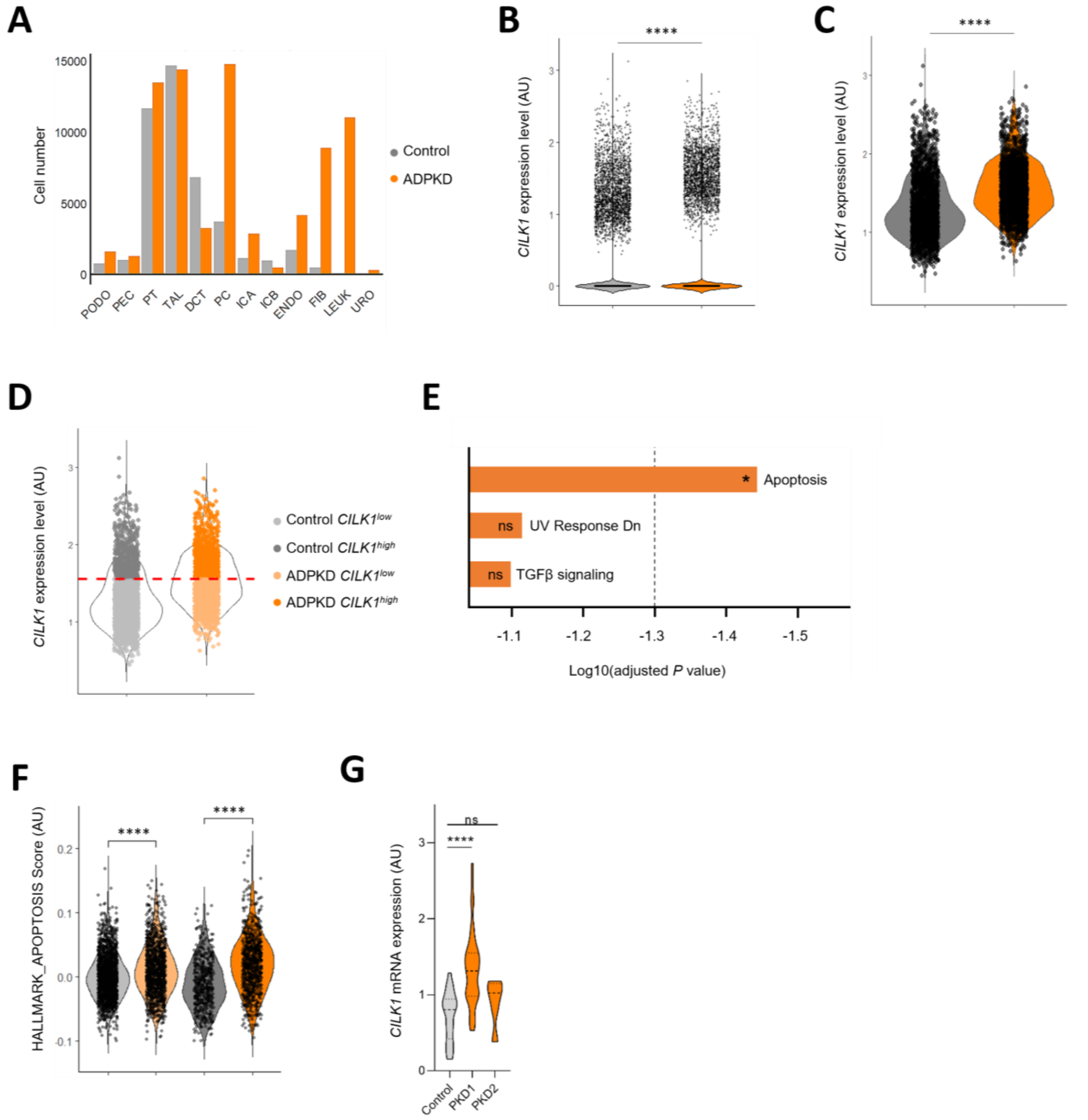


**Supplementary figure 1: ADPKD kidney cells features.** (A) Cell number per cluster. PODO podocytes; PEC parietal epithelial cells; PT proximal tubule; TAL thick ascending limb of Henle's loop; DCT distal convoluted tubule; CD collecting duct; PC principal cells; ICA Type A intercalated cells; ICB Type B intercalated cells; ENDO endothelial cells; FIB fibroblasts; LEUK leukocytes; URO uroepithelium. Histogram plot. (B) *CILK1* expression level in control (grey) and ADPKD (orange) cells. (C) *CILK1* expression level in control (grey) and ADPKD (orange) cells with a cut off expression level at 0.2. (D) *CILK1*$^{low}$ and *CILK1*$^{high}$ populations defined by the third quartile of *CILK1*-expressing control cells. (E) Top 3 up-regulated pathways (from Hallmark MSigDB) in *CILK1*$^{high}$ vs *CILK1*$^{low}$ ADPKD cells using Enrichr. (F) "Hallmark_apoptosis" (from MSigDB) score of *CILK1*$^{low}$ control (light grey), *CILK1*$^{high}$ control (dark gray), *CILK1*$^{low}$ ADPKD (light orange) and *CILK1*$^{high}$ ADPKD (dark orange) cells. (G) *CILK1* mRNA expression in controls (n=8 from Brest cohort and n=14 from Kansas cohort) and ADPKD patients (n=35 *PKD1* and n=5 *PKD2* from Brest cohort, and n=12 *PKD1* from Kansas cohort). Unpaired Student's t test:

****$P < 0.0001$, ns = non significant. (B, C, D, F, G) Violin plot. AU = arbitrary unit. (C, D, F) Cut off expression level = 0.2. (B, C, D, F) Wilcoxon test: ****P < 0.0001.

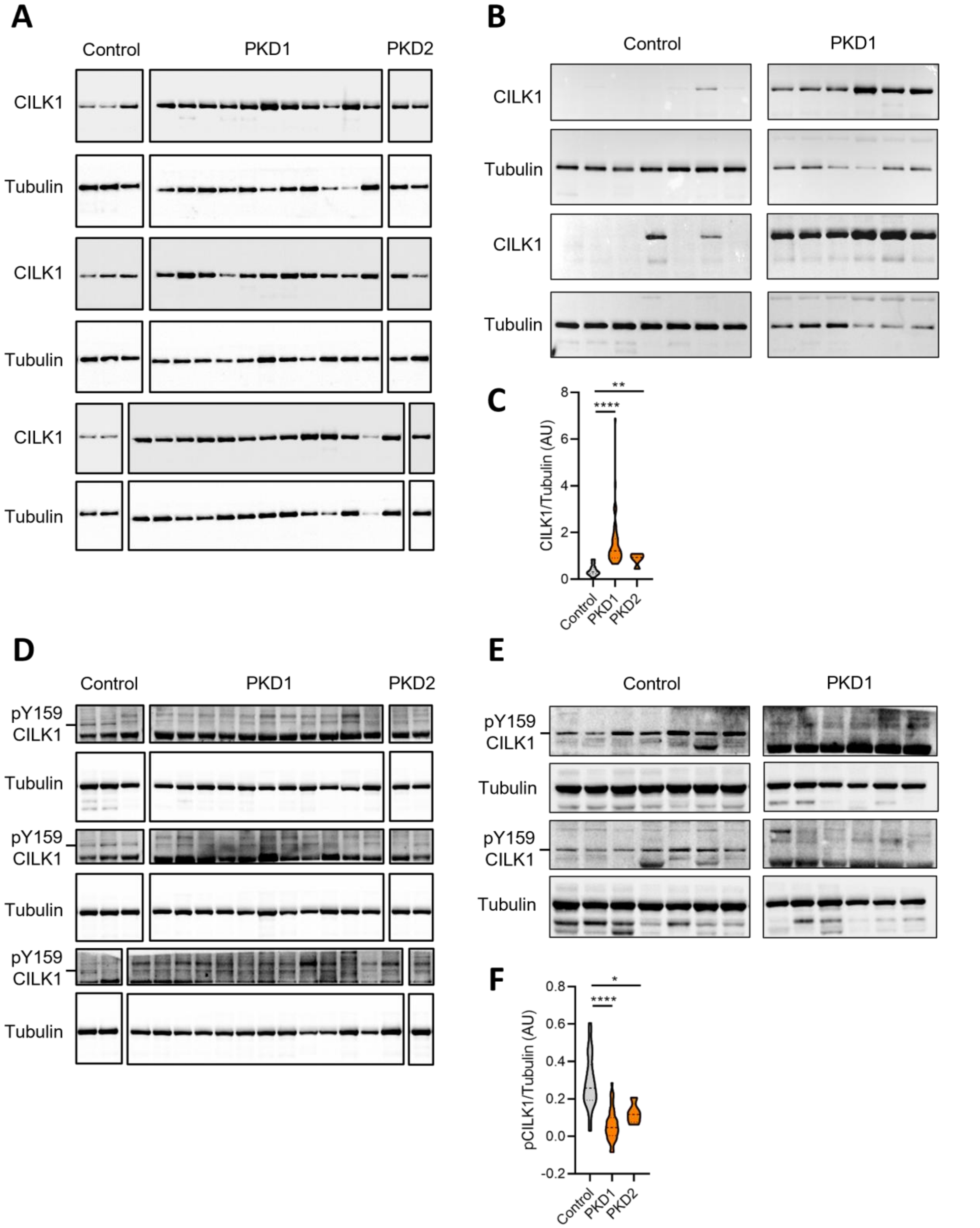


**Supplementary figure 2: CILK1 and pY159 CILK1 analysis by Western Blot.** (A, B) CILK1 Western Blot analysis of Brest cohort (n=8 controls, n=35 PKD1 and n=5 PKD2 patients; A) and Kansas cohort (n=14 controls and n=12 ADPKD patients; B). (C) Quantification of the relative abundance of CILK1. Mann-Whitney test : **$P < 0.01$, ****$P < 0.0001$. (D, E) pY159 CILK1 Western Blot analysis of Brest cohort (D) and Kansas cohort (E). (F) Quantification of the relative abundance of pY159 CILK1. Unpaired Student's t test: **$P < 0.01$, ****$P < 0.0001$.

**Supplementary tables**

**Supplementary table 1:** Transcriptional hallmarks of CILK1high cells in ADPKD samples.
**Supplementary table 2:** Patient cohorts.

**Supplementary methods**

**Cell culture**
IMCD3 cells are grown in DMEM/F12, 1% glutamine, 1% penicillin/streptomycin, 10% FBS. PKD1 guide RNAs were selected from http://crispr.mit.edu/ websites and cloned into lentiCRIPRV2 plasmids containing a blasticidin-resistance gene designed to replace the puromycin one. Lentiviruses were produced in 293FT cells after their transfection with lentiCRISPR (transfer), psPAX2 (packaging), pMD2.G (envelope) plasmids, using Lipofectamine 2000 (Life Technologies) according to the manufacturer's instructions. Supernatants containing lentiviruses were collected 48 and 72 h post-transfection. IMCD3 cells growing in a monolayer were transduced with the supernatants-containing viruses in the presence of 8 mg/mL polybrene. Transduced IMCD3 cells were selected with 6 ug/mL blasticidin (GIBCO). Oligonucleotides used for CRISPR mutations are: sgPKD1ex1#1fw CACCGGCGTCAATTGC-TCCGGCCGC, sgPKD1ex1#1rev AAACGCGGCCGGAGCAATTGACGCC, sgPKD1ex1#2fw CACCG-AGGGCCGAGTCTGCGCATCC, sgPKD1ex1#2rev AAACGGATGCGCAGACTCGGCCCTC. Cells were grown at confluency before protein extraction.

**Immunostaining**
Staining was performed on 4 µm sections. After paraffin removal and antigen retrieval treatment (10 mM Tris pH9, 1 mM EDTA, 0.05% Tween 20, 87 °C, 50 min), sections were blocked with PBS, 0.1% Tween 20, 1% BSA for 1 h and incubated overnight at 4°C with the indicated primary antibodies. After two washes, sections were incubated for 1 h with appropriate secondary antibodies and DAPI. Confocal images were acquired using a Spinning Disk microscope (10x, Andor). Areas of CILK1 staining were measured using ImageJ.

**RNA extraction and real-time RT-PCR**
Total RNA was isolated from the tissues using the RNeasy Plus kit (Qiagen) according to the manufacturer's protocol. Total RNA (50 to 1000 ng) was reverse-transcribed by the SuperScript VILOTM cDNA synthesis kit (Invitrogen) in accordance with the manufacturer's instructions. RNA expression profiles were analysed by real-time quantitative PCR using SsoFast EvaGreen Supermix (Bio-Rad) in a CFX96TM Touch real-time PCR detection system (Bio-Rad). Primers used for the detection of *CILK1* and housekeeping gene *TBP* (TATA-box binding protein) are as follows: CILK1-F 5'acaacaagcatccacagtcg3', CILK1-R 5'gaccccaccttctcctactctt3', TBP-F 5'agttgtattaacaggtgctaaa-gtcag3', TBP-R 5'gagccattacgtcgtcttcct3' (Eurofins). The complete reactions were subjected to the following program of thermal cycling: 1 cycle of 30 seconds, 50 cycles of 5 seconds at 95°C and 5 seconds at 60°C. A melting curve was run after the PCR cycles, followed by a cooling step. Each sample was run in duplicate in each experiment. Expression levels of *CILK1* were normalized to the expression level of *TBP*.

**snRNA-seq data processing**

snRNA-seq dataset of control or ADPKD tissues from the Gene Expression Omnibus (GEO) database (GSE185948) was analyzed (4). Briefly, samples were retrieved from ADPKD kidney cortical cup that were obtained from patients undergoing simultaneous native nephrectomy and living donor kidney transplantation at the University of Maryland Medical Center (Baltimore, MD). snRNA-seq on 5 control tissue (tumoral tissue from cancer patient) and 8 ADPKD tissue (cut at the base of large superficial cysts) were performed with 10X Genomics Chromium Single Cell 3' v3 chemistry. Analyses of this dataset was performed in R using Seurat (v.5.1.0). Each sample was processed as in Muto *et al (4)*, to remove low-quality nuclei (nuclei with top 5% and bottom 1% in the distribution of feature count or RNA count, or those with % mitochondrial genes > 0.25). Heterotypic doublets were identified using DoubletFinder (v.1.16.0), where the scDlbFinder score was calculated to eliminate 8% of cells with the highest threshold, as these cells are considered heterotypic doublets. After pre-processing, gene expression data for each individual sample were normalized using the *NormalizeData* function, with default parameters, which applies logarithmic normalization. For each sample, data were centered and scaled using *ScaleData* function. During scaling step, regression of S and G2M phase scores was performed to minimize the influence of cell cycle effects in downstream analysis. We used the top 2000 highly variable genes from the normalized expression matrix, to perform the principal component analysis (PCA, *runPCA* Seurat function), and applied Louvain graph-based clustering on the 30 first principal components. The resulting data were projected onto a Uniform Manifold Approximation and Projection (UMAP) using the *runUMAP* function. We applied the *FindNeighbors* and *FindClusters* functions to identify distinct clusters, using 30 dimensions, with resolution 0 to 1 with a step of 0.1. ADPKD and control kidney samples were integrated with the *IntegrateData* function with anchors identified by *FindIntegrationAnchors* function using Harmony (v.1.2.3) to create a new dimension reduction space with batch correction with *RunHarmony* function.

**Annotation of cell clusters**

For each defined cluster, a gene markers analysis was conducted using the *FindAllMarkers* function with the parameters set as follows: only.pos = TRUE, min.pct = 0.25, logfc.threshold = 0.25. The top marker genes differentially expressed selected on the average log2fold change were used to annotate cell clusters. For each cluster, we looked for known canonical markers according published data (5,6): NPHS2 and PODXL for podocytes (PODO), ALDH1A2 and CFH for parietal epithelial cells (PEC), MIOX, SLC13A3 and SLC17A1 for proximal tubule (PT), SLC12A1, UMOD and GP2 for thick ascending limb of Henle's loop (TAL), SLC12A3, KLH3 and TRPM6 for distal convoluted tubule (DCT), SCNN1G, FXDY4 and AQP2 for principal cells (PC), DMRT2, SLC26A7 and ADGRF5 for Type A intercalated cells (ICA), INSRR, SLC26A4 and SLC4A9 for Type B intercalated cells (ICB), EMCN and FLT1 for endothelial cells (ENDO), ACTA2, COL12A1 and COL1A1 for fibroblasts (FIB), PTPRC, CTSD and ZEB2 for leukocytes (LEUK), UPK1A and UPKA3 for uroepithelium (URO). Gene expression was visualized using *DotPlot* function.

**Differential gene expression and gene set enrichment analysis**

Differential gene expression analysis was performed using Seurat. To further investigate differences among epithelial CILK1-expressing cells, we classified them into two groups: CILK1-negative (*CILK1* expression <0.2) and CILK1-positive (*CILK1* expression > 0.2). To refine this analysis, CILK1-positive cells were further subdivided based on expression levels. Cells with high *CILK1* expression were defined as those above the third quartile (Q3) calculated from control patient epithelial cells, whereas cells with low *CILK1* expression were defined as those with expression levels between 0.2 and the third quartile threshold. The third quartile threshold was applied to control and ADPKD samples. Epithelial CILK1 subclusters were defined by setting cell identities with *Idents* function, and subcluster-specific marker genes were identified using *FindMarkers* function (parameters: logfc.threshold = 0.25, min.pct = 0.25). Genes with a significantly enriched p-value ($p < 0.05$, p-val > 0.25) were selected and used for pathway enrichment analysis on Enrichr website (https://maayanlab.cloud/Enrichr/). Enrichment was performed against MSigDB Hallmark 2020 database. Module scores for specific signatures (HALLMARK_MITOTIC_SPINDLE, HALLMARK_TGF, and HALLMARK_APOPTOSIS signatures, Supplementary Table 1) were calculated using the AddModuleScore function. For statistical tests comparing signature values between clusters, we used the *stat_compare_means* function and the Wilcoxon method from ggpubr R package (v.0.6.0). Data were analyzed and visualized using R (v.4.3.2). For visualization, ggplot2 package (v.3.5.1) was used and several functions from Seurat (v.5.1.0) were used such as: *ggplot*, *DotPlot*, *FeaturePlot*, *VlnPlot*.
